# Examining interactions between the microbiome and viral infection across *Drosophila* species

**DOI:** 10.64898/2026.05.27.728249

**Authors:** Ryan M. Imrie, Sarah K. Walsh, Mark A. Hanson, Xavier Harrison, Ben Longdon

## Abstract

Microbiomes can influence the outcomes of virus infection, while viral infections can disrupt microbiome composition. These reciprocal interactions may contribute to patterns of host susceptibility, but it remains unclear whether microbiome–virus interactions are consistent across host species, or whether host evolutionary relationships influence microbiome variation and its association with viral susceptibility. Here, we investigate interactions between bacterial microbiomes and Drosophila C virus (DCV) infection across 32 *Drosophilidae* host species, using experimental infections and 16S rRNA amplicon sequencing. Using phylogenetic mixed models, we tested for phylogenetic structure in microbiome diversity and composition, and whether specific bacterial taxa were associated with among-species variation in viral load. *Drosophilidae* microbiomes were consistently dominated by a small number of bacterial genera and showed limited phylogenetic structuring. DCV infection was associated with a small reduction in bacterial richness, but microbiome composition was only weakly affected by wounding or infection and largely dominated by within-species variation. However, specific taxa were associated with large differences in viral replication, including reduced DCV load in flies harbouring *Streptococcus* before infection or *Bacillus* during infection. These results suggest that, at least in the conditions used in this study, microbiome composition may not generate strong, generalisable host phylogenetic patterns in virus susceptibility.

## 1. Introduction

The outcome of viral infection can be influenced by the composition of host bacterial microbial communities, and these communities can be reciprocally disrupted by viral infection. Such bidirectional interactions have been documented across a range of systems, with commensal microbes modifying antiviral immunity, viral replication, and disease severity [1,2]. In humans, susceptibility to viral infection has been associated with variation in the bacterial gut microbiome composition [3,4], while infections with enteric and respiratory viruses – including norovirus, rotavirus, and SARS-CoV-2 – can disrupt bacterial microbial diversity and lead to normally harmless bacteria becoming pathogenic (pathobionts) [5–8]. Similar interactions can be seen in insects, where bacterial endosymbionts such as *Wolbachia* can substantially reduce replication of RNA viruses, including Drosophila C virus (DCV), Nora virus, and Flock House virus, leading to lower viral titres and increased host survival [9,10]. These effects are variable, and in some systems *Wolbachia* has been associated with increased susceptibility to viral infection [11,12]. Reciprocally, viral infections in insects can alter bacterial load and the activation of immune pathways associated with host-bacteria interactions [13–15].

Most evidence for interactions between viruses and the microbiome (hereafter microbiome is used to refer to the bacterial community within a host) has come from studies of individual host species. However, variation in both virus susceptibility and microbiome composition are frequently studied separately in comparative analyses across host species. Studies of phylosymbiosis – the association between microbiome community composition and host evolutionary relationships – have revealed that microbiome composition is often correlated with host phylogeny across a wide range of different animal taxa [16]. These patterns are particularly strong in vertebrates and mammals, where closely related species tend to harbour more similar microbial communities [17]. In insects, evidence for phylosymbiosis is more mixed. While some studies have identified phylogenetic signals in gut bacterial communities across taxa such as ants, parasitoid wasps, and social bees [18–20], others – including studies of *Drosophila* and *Coleoptera* – find weaker or inconsistent associations [21–24]. In *Drosophila*, the gut microbiome is often transient and environmentally acquired [25], and the inconsistent evidence of phylosymbiosis observed in these hosts may partly reflect the loss of species-specific microbial exposure under laboratory conditions [21]. Yet, studies of wild-derived bacterial strains and fly lines have shown that stable and host-specific microbiome associations can occur in *Drosophila* [26,27]. Lab-reared flies typically harbour low-diversity communities dominated by Acetobacter and Lactobacillus (recently reclassified into 25 genera within the Lactobacillaceae family [28]). The simplicity of the lab-reared fruit fly microbiome has made Drosophila melanogaster a widely used model for mechanistic studies of host–microbiome interactions [29].

Comparative analyses of virus susceptibility across host species have consistently shown that large proportions of variation of virulence, transmissibility, and virus replication can be explained by the evolutionary relationships between hosts – with phylogeny acting as a proxy for divergence in underlying traits (e.g., physiology and immunity) across hosts [30]. Studies of vertebrate viruses have shown that host species more distantly related to humans harbour fewer zoonotic viruses [31], and that viruses transmitted from these hosts tend to show higher case fatality rates and lower human-to-human transmissibility [32]. Similar patterns were seen in studies of rabies cross-species transmission, where host shifts become less frequent with increasing phylogenetic distance between hosts [33], and spillover infections from more distantly related hosts tend to have shorter clinical periods, reducing opportunities for onward transmission [34]. In *Drosophila*, variation in virulence during both bacterial and viral infection can be explained by the relationships between hosts [35–37], with more closely related species tending to experience more similar infection phenotypes. Similar patterns are seen when examining variation in viral load, with the *Drosophila* phylogeny explaining a large proportion of across species variation for both RNA and DNA viruses, although viral loads are only weakly correlated across virus families, consistent with host-by-virus interactions in infection outcome [38,39]. Feasibly, interspecific variation in virus susceptibility may be partly due to variation in the microbiome composition across host species, while more susceptible hosts may in turn provide greater opportunity for virus infection to perturb the resident microbial community; however, these possibilities remain largely unexplored [40].

Mechanistically, reciprocal virus-microbiome interactions may occur for a number of reasons. In vertebrates, microbiota can inhibit viral infection through immune modulation or direct antagonism, including induction of type I interferon [41,42] and direct virucidal effects such as the lactic acid secreted by *Lactobacillus* [43]. Conversely, microbiota can enhance infection, for example by stabilising virions or facilitating host cell attachment [44,45], and viral infections can perturb the microbiome through tissue damage, altered nutrient availability, or systemic immune activation [1]. Both respiratory and enteric viral infections can disrupt the balance of host epithelial immunity and the microbiome, reducing gut microbial diversity and promoting the expansion of pathobionts through inflammation and loss of mucosal integrity [22,46]. In insects, similar immunological interactions can be seen in the priming of antiviral pathways by gut commensals [47], and virus-induced changes in gut homeostasis and integrity [48,49]. In *Drosophila*, Gram-negative bacteria such as *Acetobacter* activate the Imd pathway via peptidoglycan recognition, promoting downstream ERK signalling and restricting enteric viral replication (including DCV) [50,51]. Reciprocally, sensing of the DCV suppressor of antiviral RNAi induces the lncRNA VINR, which activates a non-canonical antimicrobial pathway and promotes antimicrobial peptide expression, and loss of VINR increases susceptibility to both DCV and Gram-negative and Gram-positive bacterial infection [52].

It remains unclear the extent to which virus-microbiome interactions are dependent on host context, or whether interactions are broadly similar across host species. In insects, the endosymbiont *Wolbachia* is thought to reduce viral replication through metabolic competition and host cell reprogramming, creating intracellular environments that are unfavourable for viral growth [53]. This interaction appears to be conserved across Dipteran hosts, having been studied extensively in *Drosophila* [9,54–56] and applied translationally to wild mosquitoes [57,58]. However, *Wolbachia*-mediated protection is not universal, with some instances where *Wolbachia* can increase viral susceptibility [11,59]. *Wolbachia* load varies across host species [60], and the protection phenotype appears to be driven primarily by *Wolbachia* strain effects linked to bacterial load [61,62]. More generally, if host genetic background influences microbial composition, viral replication, or the host pathways that mediate their interaction, then virus-microbiome dynamics may show detectable phylogenetic structure.

Here, we investigate how bacterial microbiomes interact with RNA virus infection across host species, using experimental infection with Drosophila C virus (DCV) across a panel of 32 *Drosophilidae* host species. Combining 16S rRNA amplicon sequencing with phylogenetic mixed models, we compare microbiome diversity and composition in uninjected (control), saline injected, and DCV infected flies, and test for phylogenetic structure in these microbial communities. We then quantify whether variation in bacterial taxa across host species is associated with differences in viral replication, allowing us to assess the extent to which microbiome composition contributes to interspecific variation in virus susceptibility, and whether virus–microbiome interactions show consistent patterns across the *Drosophilidae* phylogeny.

## 2. Materials & Methods

### 2.1 Fly Species

Laboratory stocks of 32 *Drosophilidae* host species (Supplementary Table 1) were maintained in multigeneration stock bottles at 22°C under a 12-hour light-dark cycle. Each bottle contained 50ml of one of four media varieties, selected to optimize rearing conditions (recipes available at doi:10.6084/m9.figshare.21590724.v1). Although these diets differ in macronutrient composition, variation in laboratory media has been shown to have minimal influence on the outcome of viral infection in this system [63]. To account for the possibility of diet affecting the reciprocal interactions between the microbiome and DCV, rearing diet is included as a random effect in all models (see below for details). Note that during experiments, adult flies were kept on a single diet type, with repeated tipping, so had similar environmental exposure to microbes, which has previously been reported to lead to similar microbiomes across lines/species [25,29,64,65].

The host phylogeny used in this study was inferred in BEAST v1.10.4 [66] following steps described previously [37]. Briefly, publicly available sequences for the *28S, Adh, Amyrel, COI, COII, RPL32*, and *SOD* genes were obtained from GenBank (see https://doi.org/10.6084/m9.figshare.13079366.v1 for a full list of accessions by species). Sequences were aligned in Geneious v9.1.8 (https://www.geneious.com/) using progressive pairwise global alignment with free end gaps and a 70% similarity IUB cost matrix. Default settings were used for gap opening, gap extension, and refinement iterations.

Phylogenetic reconstruction was performed using a relaxed clock model in BEAST, as the phylogenetic mixed models used for the statistical analysis of microbiome interactions require a tree with equal root-to-tip distances across taxa. Genes were partitioned into ribosomal (*28S*), mitochondrial (*COI, COII*), and nuclear (*Adh, Amyrel, RPL32, SOD*) groups. The mitochondrial and nuclear partitions were further divided by codon positions (1 + 2 and 3), with unlinked substitution rates and base frequencies across partitions. Each partition was fitted to a relaxed uncorrelated lognormal molecular clock model using random starting trees and four-category gamma-distributed HKY substitution models. Two independent BEAST runs were conducted with 1 billion MCMC generations each, sampling every 100,000 iterations with a birth-death process prior on the tree shape.

Model trace files were assessed for convergence, sampling efficiency and autocorrelation in Tracer v1.7.1 [67]. A maximum clade credibility tree (Supplementary Figure 1) was inferred from the posterior distribution after a 10% burn-in, showing topologies broadly consistent with *Drosophilidae* phylogenies based on larger gene sets [68].

### 2.2 Inoculation

The isolate of DCV used in this study (DCV-Charolles; DCV-C) was kindly provided by Julien Martinez [69] and was previously verified to be free of contamination with other common *Drosophila* viruses via qRT-PCR [38]. Before inoculation, 0-1-day-old male flies were collected and maintained in vials containing cornmeal media at 22°C and 70% relative humidity under a 12-hour light-dark cycle. Each vial held between 8 and 12 flies (mean = 11.1). Flies were transferred to fresh media every two days until they reached 7-8 days of age, when they were anesthetized with CO_2_ and either a) left anesthetized for 5 minutes (“no injection”); b) inoculated with Ringers (“saline injection”); or c) inoculated with DCV (“DCV infection”) at a TCID50 of 6.32×10^9^. Inoculations were performed by septic pinprick using a 12.5μm diameter stainless steel needle (Fine Science Tools, CA, USA). Each needle was bent to a right angle approximately 250μm from the tip to serve as a depth stop and inserted into the right lateral anespisternal cleft up to the angle, to provide a consistent inoculation depth. In total, 3,184 flies were inoculated across the experiment, organised into three biological replicate blocks (vials) per combination of host species and inoculation condition (see Supplementary Table 2 for a breakdown of replicates across species and treatments).

This inoculation method bypasses the gut immune barrier but avoids dosage variation associated with differences in feeding behaviour among host species [70]. Only males were used to eliminate potential confounding effects of sex or mating status, both of which can introduce additional variation in pathogen susceptibility [71–73]. Previous work has shown strong positive correlations between male and female DCV susceptibility in these host species [74].

### 2.3 RNA and DNA Extraction

After 2 additional days (+/-2 hours), flies were snap frozen in liquid nitrogen then homogenized in Trizol using an Omni Bead Ruptor 24 (4 m/s for 15 seconds) with 0.2mm Zirconia beans (Thistle Scientific). The 2 dpi timepoint was selected based on previous work which showed this period was sufficient for interspecific variation in viral load to appear but not so long that infection-related mortality reduced within-vial sample sizes [37]. All samples underwent a dual RNA-DNA extraction process. First, total RNA was extracted using a standard Trizol chloroform-isopropanol extraction method. The remaining organic and lower aqueous phases were then mixed with a back extraction buffer (4M guanidine thiocyanate, 50mM trisodium citrate, 1M TRIS), producing a DNA-containing upper phase for collection. Isolated DNA was precipitated overnight in isopropanol at -20C, pelleted, and washed three times with 70% ethanol before resuspension in 8mM NaOH. After resuspension, DNA samples were pH neutralised with 1M HEPES.

### 2.4 Viral Load Measurement

qRT-PCR was carried out for DCV and the housekeeping gene *RPL32* using the Sensifast Lo-Rox SYBR kit (Bioline) on an Applied Biosystems QuantStudio 3 (primer sequences and cycling conditions are provided in Supplementary Tables 3-5). Between-plate variation in Ct values was corrected for statistically using standard methods [75,76]. Product specificity was confirmed by melt curve analysis, using ±1.5 °C and ±3 °C inclusion thresholds for DCV and *RPL32* amplicons respectively, and two technical replicates per sample were collected and averaged to obtain mean Ct values for DCV and *RPL32*.

To estimate the inoculation dose, eight vials of *D. melanogaster* (20 males per vial) were snap frozen immediately after DCV inoculation and processed as above. These values were used to infer the RPL32-normalized inoculation of other host species as:

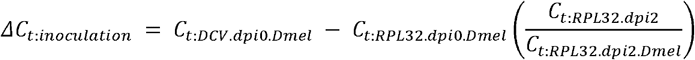

Applying this approach to data from a previous study in which inoculation doses were measured separately for each species showed that the method accurately inferred normalized doses across species (R^2^ = 0.92) [38]. Fold changes in viral load for each biological replicate were then calculated using the 2^−^ΔΔCt method as *ΔΔC*_*t*_ *=ΔC*_*t:inoculation*_ − *ΔC*_*t:dpi2*_.

### 2.5 16S Sequencing

Bacterial *16S* ribosomal RNA (rRNA) gene V3-V4 regions were amplified by conventional PCR (for cycle conditions see Supplementary Table 5) using the prokaryote universal primers 16S_341F (5’-CCTACGGGNGGCWGCAG-3’) and 16S_785R (5’-GGACTACHVGGGTATCTAATCC-3’). PCR clean-up was performed using AMPure XP Beads (Beckman Coulter), and amplicon libraries generated with the Nextera XT DNA Library Preparation kit (Illumina). Indexed libraries were purified with AMPure XP beads and eluted in 10 mM Tris pH 8.5. Library size was verified by Bioanalyzer (Agilent DNA 1000 kit), with the expected final product length of ∼630bp. Library concentrations were measured using a Qubit dsDNA BR Assay kit (Invitrogen) and normalised to 4nM before pooling. Sequencing was performed on an Illumina MiSeq using a 300bp paired-read v3 600-cycle kit by the Exeter Sequencing Service. The following controls were included in the sequencing run in addition to experimental samples: 6x negative extraction controls (nuclease-free water), 6x Microbial Community Standards (ZymoBIOMICS) and 6x Microbial Community DNA Standards (ZymoBIOMICS). Each control sample was spiked into Trizol and underwent RNA/DNA extraction, 16S PCR amplification, and library preparation in tandem with experimental samples.

### 2.6 Bioinformatics

Raw 16S reads were processed using Cutadapt [77] to remove adaptor sequences, after which all subsequent quality filtering, trimming, denoising, and error correction were performed in R using the dada2 v1.26 pipeline [78]. Reads were truncated to 240 bp (forward) and 225 bp (reverse) and filtered to retain only those with ≤2 expected sequencing errors. Error rates were learned independently for each read direction, and denoised reads were used to infer unique amplicon sequence variants (ASVs). ASVs were retained if their merged length matched the expected size range for the V3–V4 region and were assigned taxonomy using the SILVA v138.1 reference database [79]. To infer an ASV phylogeny, unique V3–V4 sequences were aligned with MAFFT [80], low-information regions were removed with trimAL [81], and an approximate maximum-likelihood tree was generated using FastTree 2 [82].

Contaminants were removed in several steps: non-bacterial sequences (chloroplasts and archaea) were manually pruned, and technical contaminants were identified using a prevalence-based approach in decontam [83]. Additionally, ASVs were removed if their mean abundance in experimental samples did not exceed their mean abundance in negative controls, resulting in the removal of 3 ASVs. Samples were filtered to remove those with fewer than 1,350 reads remaining after processing. This cut-off was chosen as the highest value that prevented any combinations of host species and infection condition from falling below two biological replicates. Finally, low-prevalence taxa were filtered by retaining only ASVs present at ≥1 copy in at least 5% of remaining biological replicates [84], resulting in the removal of 285 ASVs. This produced a final dataset containing 50 ASVs, all of which were resolved to the genus level, and 11 of which resolved to the species level (Supplementary Table 6).

To provide an ultrametric phylogeny for the final 50 bacterial ASVs, full-length 16S rRNA sequences from the SILVA v138.1 database were nucleotide-aligned using MUSCLE 5.1 [85] with default settings. The alignment was used to fit a relaxed uncorrelated lognormal molecular clock model in BEAST using a random starting tree, a four-category gamma-distributed HKY substitution model, and no codon partitioning. Model runs, trace file validation, and the production of a maximum clade credibility tree were performed identically to the *Drosophilidae* phylogenetic inference (see above).

### 2.7 Statistical Modelling

All statistical analyses were performed in R [86] using the MCMCglmm package [87]. As 16S sequence data are compositional in nature, centre log ratio (CLR)–transformed abundances were used for beta diversity and differential abundance analyses, calculated using a count-zero-multiplicative (CZM) approach [88,89], whereas alpha diversity metrics were calculated directly from raw read counts.. Alpha diversity metrics (richness, evenness, and Shannon diversity) were calculated using the phyloseq package [90]; and beta diversity explored by principal component analysis (PCA, stats package) and non-metric multidimensional scaling with an Aitchison distance metric (NMDS, vegan package). Due to the zero-inflated nature of the 16S data, models using CLR-transformed abundance as a response were fitted as multivariate models combining binary presence/absence (determined from raw reads) modelled as a threshold trait with a probit link, and continuous CLR-transformed abundances modelled with a Gaussian error distribution. Random effects within each model followed one of two general structures: a structure intended to estimate phylogenetic heritability (1); and a structure intended to estimate repeatability (2):

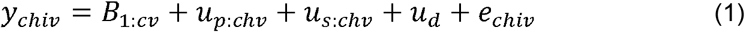

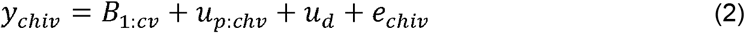

In these models, *y*_*chig*_ is the response variable for the combination of injection condition *c* (uninjected, saline injected, DCV infected), in the *ith* biological replicate of host species *h* and bacterial ASV *v*. The fixed effect *B*_1_ is the intercept for each injection condition; the random effect *u*_*p*_ represents the effect of host phylogeny under a Brownian model of evolution; *u*_*s*_ represents a categorical host-species-specific effect that is independent of phylogeny and explicitly estimates the non-phylogenetic component of variance. Lastly, *u*_*d*_ represents a random effect of rearing diet, and *e* is the model residuals, both of which also assumed Gaussian errors. Credible intervals for the proportion of total variance attributable to rearing diet included zero in all models, but also frequently spanned non-zero values (Supplementary Table 8) and so rearing diet was retained in all models. For response variables that summarise across bacterial ASVs (e.g., alpha/beta diversities) *v* is absent from the model structure.

The proportion of among-host-species variation that can be explained by host phylogeny (equivalent to phylogenetic heritability or Pagel’s lambda) was calculated as 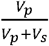, where *v*_*p*_ and *v*_*s*_ are the phylogenetic and species-specific components of variance respectively, taken from model structure (1). The repeatability of measurements within-host-species after accounting for any phylogenetic variance was calculated as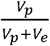, where *v*_*p*_ and *v*_*e*_ are the phylogenetic and residual variance components from model structure (2). In multivariate models, covariance matrices were used to calculate correlation coefficients between conditions as 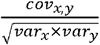 and regression slopes as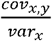. Intercepts for these correlations were calculated as 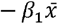, where *β*_1_ is the fixed-effect estimate of the across-species mean. For marginal correlations, variance components were summed across matrices as *v*_*p*_ +*v*_*s*_ +*v*_*e*_ before calculating correlation coefficients. For parameters on an unbounded scale (correlation coefficients, slopes), effects were considered significant when the 95% credible intervals (calculated here as 95% HPD intervals) did not include zero. For proportions of variance (heritability, repeatability), effects were considered significantly non-zero when the lower bound of the 95% credible interval exceeded a value of 0.05. This threshold was chosen to avoid treating effects whose posterior distributions allocate a high probability mass near zero as meaningful.

Each model was run for 13,000,000 MCMC generations, sampled every 5,000 generations after a burn-in of 3,000,000 generations. Presented results are taken from models with parameter-expanded priors placed on the phylogenetic (*v*_*p*_) and species-specific (*v*_*s*_) covariance matrices, and inverse-gamma priors placed on the residual variances. Alternative models were also fitted with flat and inverse-Wishart priors, which provided qualitatively similar results. Model convergence was assessed by manually inspecting parameter posterior distributions, effective sample sizes (ESSs) and autocorrelation diagnostics. Full details on the structure of each individual model, and all scripts and data used in this analysis are available at https://github.com/ryanmimrie/Publications-2026-Drosophilidae-16S-Virus-Infection.

## 3. Results

To investigate the variation existing in the bacterial microbiome across 32 species of lab-reared *Drosophilidae*, and how such variation may influence or be influenced by RNA virus infection, microbiome compositions from each species were characterised by 16S amplicon sequencing following either DCV infection, saline injection, or no treatment controls (referred to here as “uninjected”). Bacterial amplicon sequence variants (ASVs) varied in relative abundance among host species and experimental conditions (Figure 1), with microbiomes consistently dominated by a small number of genera: *Commensalibacter, Acetobacter, Serratia, Pseudomonas*, and *Dysgonomonas* together accounted for 86% of all sequenced bacterial reads.

**Figure 1:**
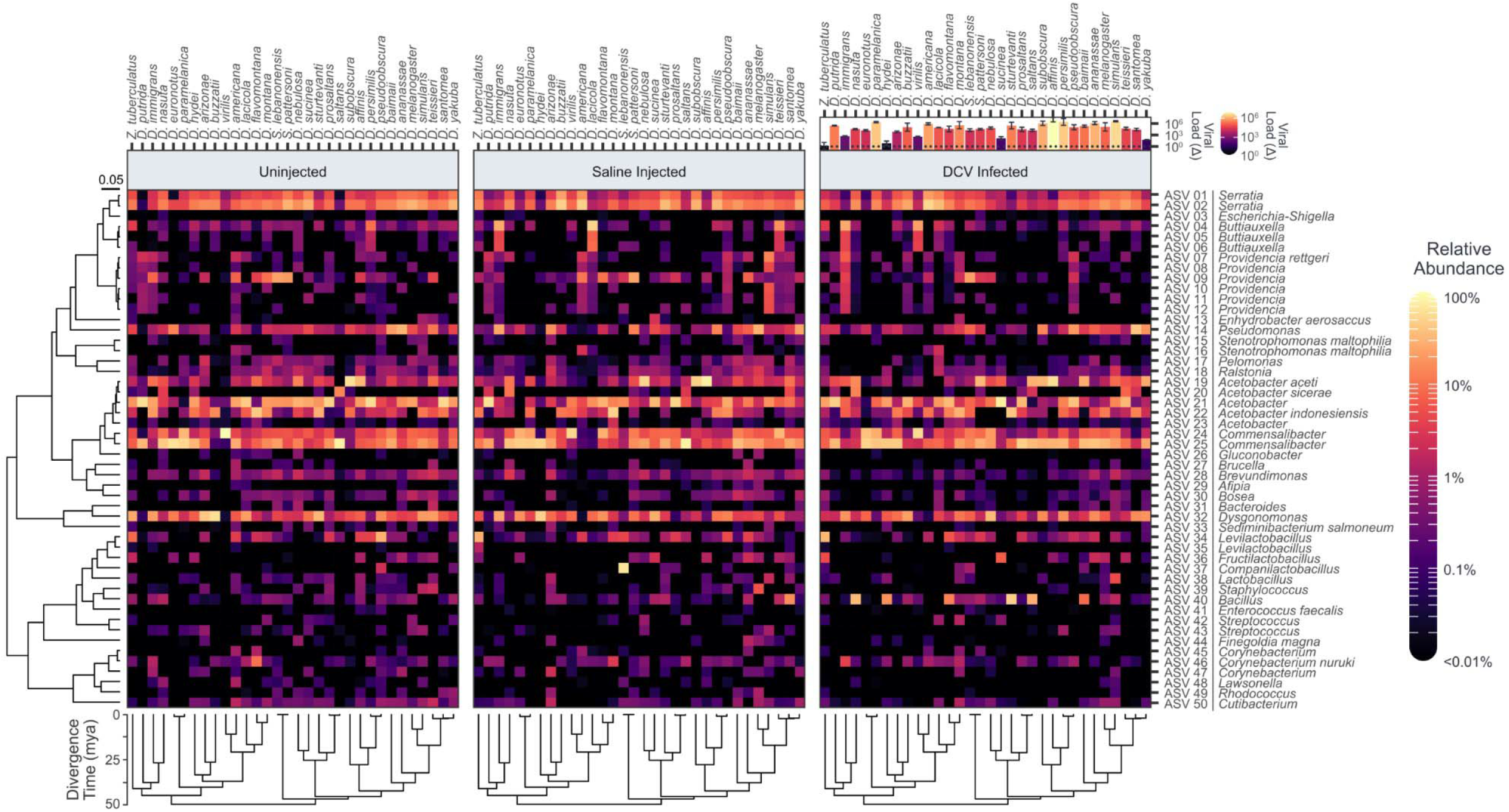
Relative abundances of bacterial ASVs across *Drosophilidae* host species and injection conditions. Heatmaps show the mean relative abundance of bacterial ASVs (rows) in different host species (columns) on a log_10_ scale, separated by injection condition. Maximum clade credibility phylogenies are presented for both bacteria (left) and *Drosophilidae* (bottom) with a scale bar for the number of nucleotide substitutions per site. For the *Drosophilidae* phylogeny, the approximate age since divergence, based on estimates from [100] is shown in the axis at the bottom left. Above the DCV infection panel, bar height and colour show the mean change in viral load from 0 to 2 dpi on a log_10_ scale, with error bars representing the standard error of the mean.

DCV infection resulted in a small but significant reduction in bacterial ASV richness across host species, with an average of 2.1 fewer ASVs (95% CI: –3.9, –0.14) detected compared to uninjected flies, which had a mean richness of 17.1 ASVs (95% CI: 14.9, 19.3). In contrast, saline-injected flies did not show a detectable change in richness relative to uninjected controls (mean difference = -0.52, 95% CI: -2.8, 1.6), indicating that the reduction in richness was associated with viral infection rather than wounding alone. Other measures of alpha diversity (evenness and Shannon diversity) showed no significant differences between experimental conditions (Figure 2A). When analysed using phylogenetic GLMMs, saline-injected flies yielded significantly non-zero estimates of phylogenetic heritability and repeatability for all three alpha diversity metrics (Figure 2B), while DCV-infected flies showed significantly non-zero estimates for heritability and repeatability of bacterial ASV evenness and Shannon diversity. No significant estimates of heritability or repeatability were detected for any alpha diversity metric in uninjected flies. Analyses of beta diversity using principal-component analysis (PCA, Supplementary Figure 3A) and non-metric multidimensional scaling (NMDS, Supplementary Figure 4A) show little tendency for microbiome compositions to cluster by experimental condition. Correspondingly, PERMANOVA detected only a very small but consistent effect of experimental condition on CLR-based β-diversity (R^2^ = 1.25%, F_2,265_ = 1.68, p = 0.019), which remained detectable when permutations were constrained within host species (although note that these analyses do not take into account the non-independence of species due to common ancestry). Individual axes of both the PCA and NMDS showed evidence of phylogenetic heritability and repeatability, including in uninjected flies (Supplementary Figure 3B, Supplementary Figure 4B). Together, these results suggest that wounding and DCV infection introduce subtle shifts in microbiome diversity across host species and, independent of experimental condition, microbiome composition shows limited evidence of structuring by host phylogeny along specific multivariate axes.

**Figure 2:**
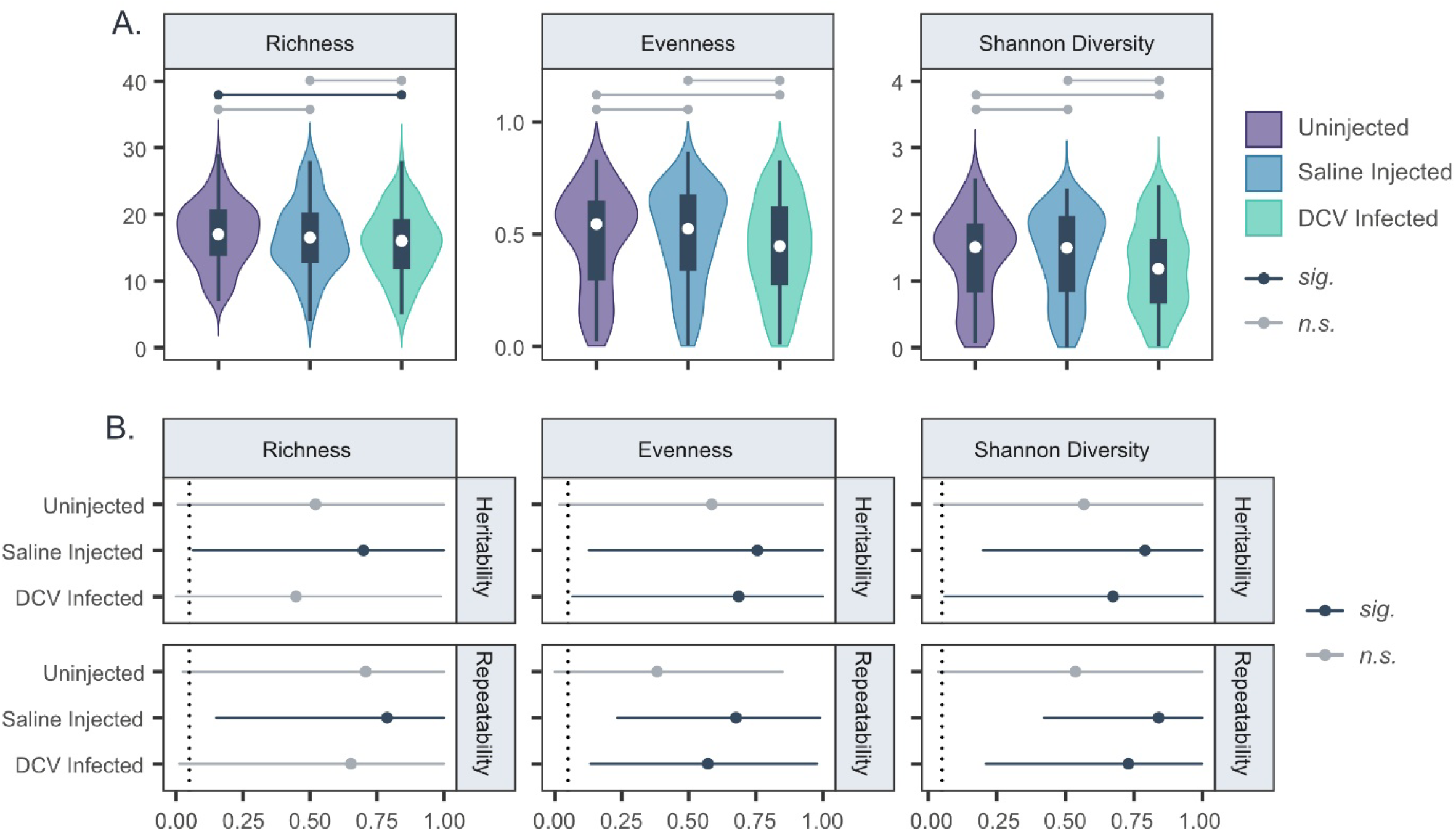
Alpha Diversity Metrics. A) Violin plots of the distribution of alpha diversity metrics across-host species and experimental conditions, showing richness (left facet), evenness (middle facet) and Shannon diversity (right facet). B) Estimates of phylogenetic heritability (top facet row) and repeatability (bottom facet row) for each alpha diversity metric under each experimental condition. Points represent the mean and bars the 95% CIs of the posterior distributions of each estimate. Significant differences between conditions (A) and non-zero estimates (B) are highlighted in black.

As many bacterial ASVs were absent from multiple samples, patterns in individual genera across host species were analysed using hurdle models [91], multivariate phylogenetic GLMMs that estimate a detection probability from the binary presence/absence component of the 16S data (Figure 3A) and a continuous measure of abundance in samples where the genus is present (Figure 3B). Across these models, variation in presence and abundance consistently failed to segregate into phylogenetic or species-specific random effects (Supplementary Table 7-8), suggesting that much of the variation in individual bacterial ASVs occurs within *Drosophilidae* host species. Four ASVs showed significant decreases in detection probability following saline injection (ASV 04: *Buttiauxella, ASV 14: Pseudomonas, ASV 43: Streptococcus*, and ASV 50: *Cutibacterium*), and one ASV increased in detection probability (ASV 09: *Providencia*). Additionally, four ASVs showed decreases in detection probability in DCV infected flies compared to saline injected flies (ASV 17: *Pelomonas*, ASV 18: *Ralstonia*, ASV 19: *Acetobacter aceti*, and *ASV 47: Corynebacterium*), with no ASVs showing a credible increase during infection. Two ASVs showed detectable changes in abundance with saline injection, one increasing (ASV 04: *Buttiauxella*) and another decreasing (ASV 31: *Bacteroides*). Lastly, five ASVs showed significant decreases in abundance with DCV infection (ASV 17: *Pelomonas*, ASV 18: *Ralstonia*, ASV 28: *Brevundimonas*, ASV 30: *Bosea*, and ASV 37: *Companilactobacillus*), and two increased in abundance (ASV 19: *Acetobacter aceti* and ASV 40: *Bacillus*). For most ASVs, no significant changes in detection probability or abundance were associated with wounding or infection, leading to strong marginal correlations across infection conditions (Figure 4).

**Figure 3:**
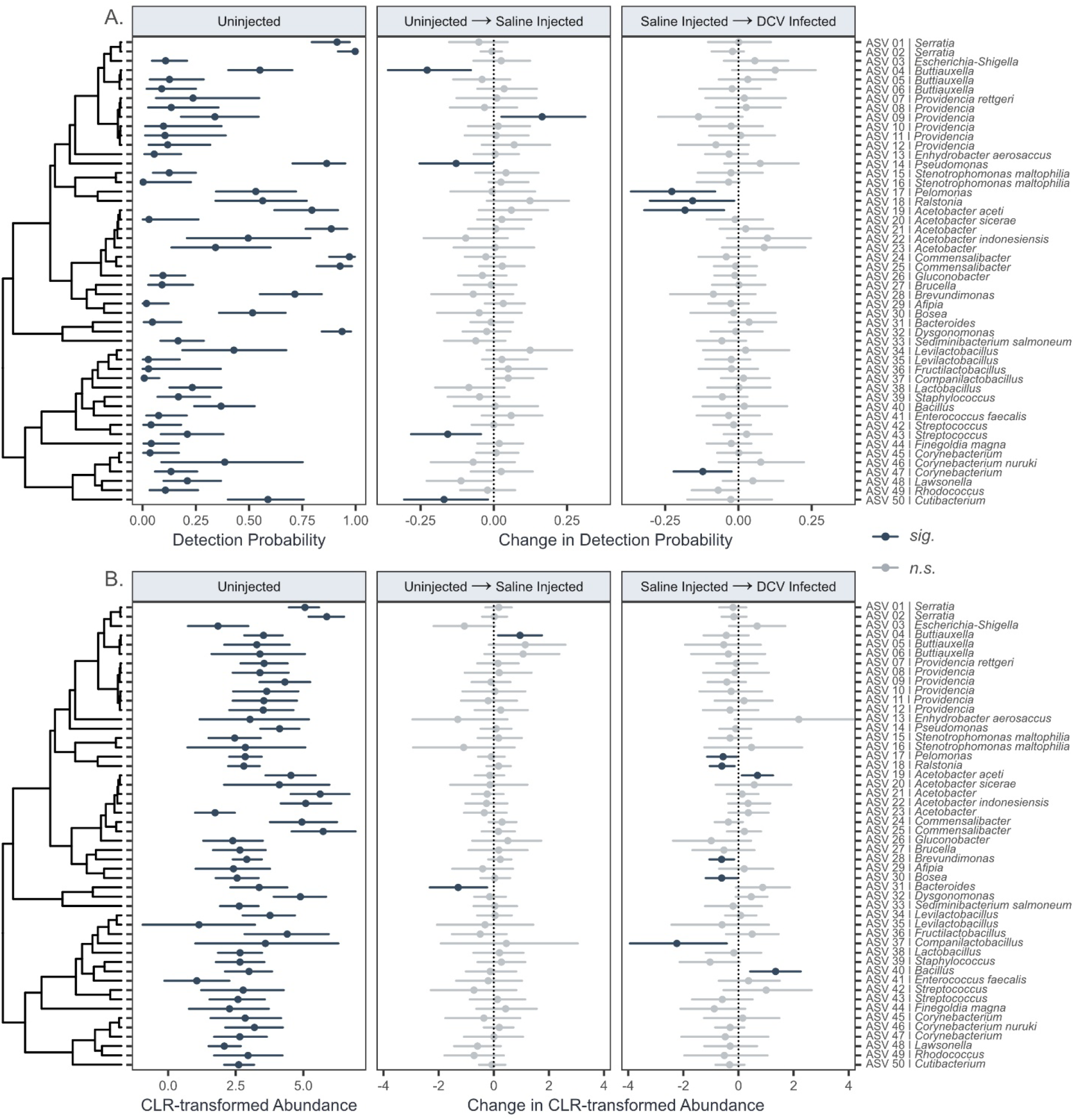
Changes in mean presence and abundance of bacterial ASVs with injection and infection. Values for the across-host-species detection probability (A) and Centre Log Ratio (CLR) transformed abundance (B) in uninjected flies (left facet) and the change in detection probability following septic injury (middle facet) and DCV infection (right facet). Points represent the mean and bars the 95% HPD intervals of the posterior distribution of each estimate taken from a model of structure (2). Estimates of the change in detection probability or CLR-transformed abundance that do not include zero are highlighted in black.

**Figure 4:**
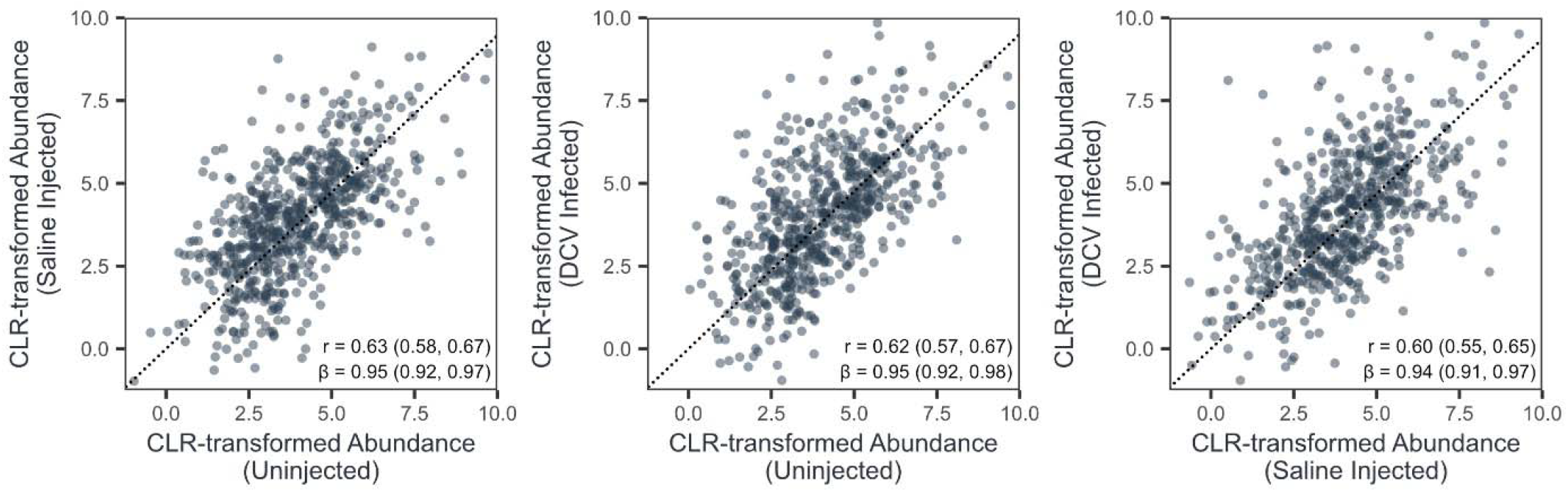
Correlations in abundance of bacterial ASVs between injection conditions. Correlations in Centre Log Ratio (CLR) transformed abundance between uninjected and saline injected flies (A); uninjected and DCV infected flies (B); and saline injected and DCV infected flies (C). Individual points represent the mean abundance for each *Drosophilidae* host species and bacterial ASV. Marginal correlation coefficients (r) and trendlines are taken from a phylogenetic mixed model of structure (1) as phylogenetic and species-specific covariance matrices from models of structure (1) and (2) contained very small (<0.01) proportions of total variance in bacterial abundance.

Feasibly, microbial communities may affect the outcome of RNA virus infection due to conditions that exist before infection (considered here to be represented by the uninjected fly condition), or during infection, and these effects may manifest at the community level (alpha and beta diversities), or at the level of individual ASVs. No significant correlations were detected between DCV viral load and any alpha diversity metrics across host species. Similarly, no PC or NMDS axis showed detectable interspecific correlations with DCV viral load (Supplementary Table 9). Additionally, two significant marginal correlations between DCV viral load and individual bacterial ASV presence were detected across host species (Figure 5, Supplementary Figure 5A). Fly species with no detectable *Streptococcus* in uninjected flies had a mean fold-change in viral load of 4.0 × 10^4^(95% CI: 1.6 × 10^3^, 8.8 × 10^5^). In host species where *Streptococcus* was present in uninjected flies, the estimated mean fold-change in viral load was ∼4.4 × 10^3^, with the posterior contrast indicating an 89% reduction in viral load (95% CI: 70%, 96%). Similarly, fly species lacking detectable *Bacillus* during infection had a mean fold-change in viral load of 3.9 × 10^4^ (95% CI: 1.4 × 10^3^, 1.26 × 10^6^). When *Bacillus* was present the mean fold-change in viral load was ∼3.9 × 10, with the posterior contrast indicating a 90% reduction in viral load (95% CI: 69%, 96%). No significant marginal correlations between DCV viral load and bacterial ASV abundance were detected across host species (Supplementary Figure 5B).

**Figure 5:**
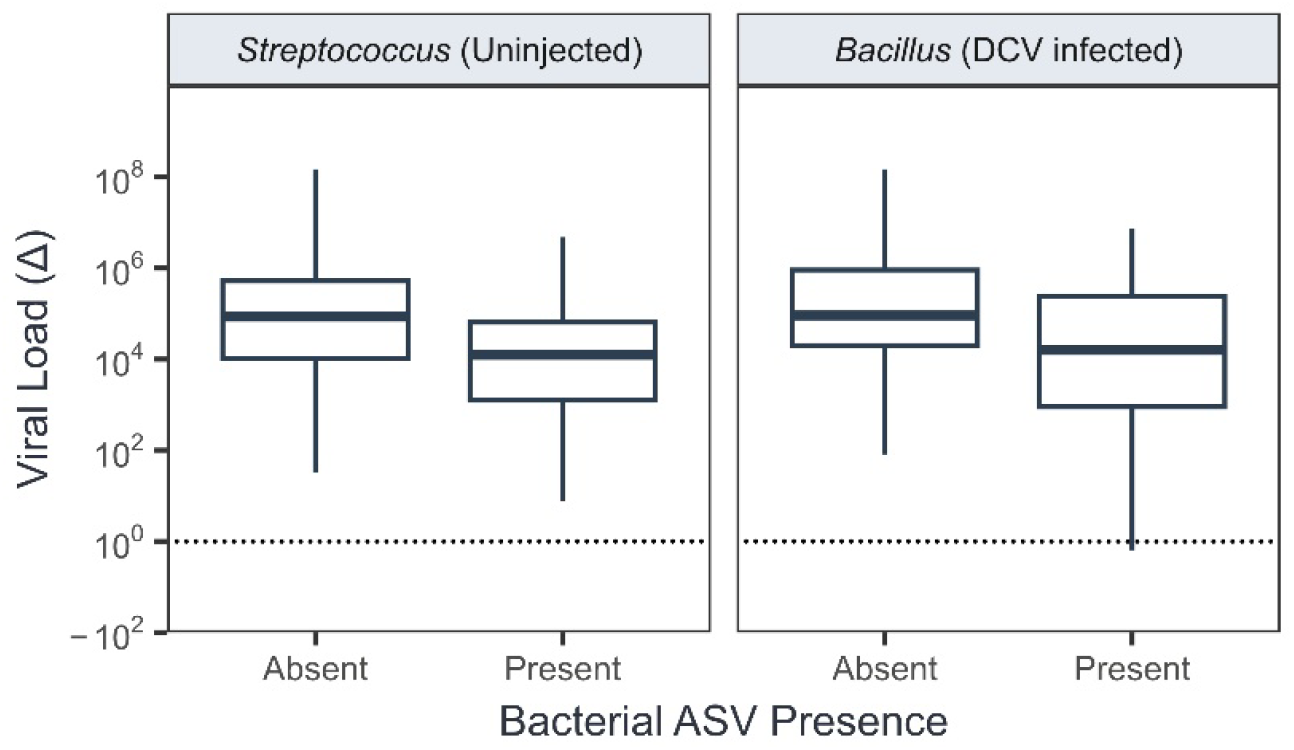
Significant correlations between viral load and bacterial ASV presence before (uninjected) and during (DCV infected) infection. Boxplots show the relationships between fold-change in DCV viral load from 0-2 dpi on a log_10_ scale and the detectable presence of *Streptococcus* before infection (left facet) and *Bacillus* during infection (right facet).

## 4. Discussion

Here, we examined interactions between bacterial microbiomes and RNA virus infection across 32 *Drosophilidae* host species, using 16S rRNA amplicon sequencing and experimental infection with DCV. Across host species, microbiomes were consistently dominated by a small number of bacterial genera, and community composition showed limited evidence of host phylogenetic structuring. Septic injury and DCV infection caused only subtle shifts in community diversity and composition, including a small reduction in microbiome richness during infection. Despite this apparent stability at the community level, DCV infection was associated with among species changes in the presence and abundance of a small number of bacterial taxa. In turn, the presence of two bacterial taxa were associated with a general reduction in DCV replication. Together, these results align with previous work indicating that *Drosophila* microbiomes show weak and inconsistent evidence of phylosymbiosis in laboratory settings [22,24]. However, we show that virus–microbiome interactions in *Drosophilidae* include a small number of bacterial taxa whose interactions with virus infection are common and detectable across hosts.

In response to DCV infection, four bacterial ASVs showed a reduction in detection probability (*Ralstonia, Pelomonas, Acetobacter aceti*, and *Corynebacterium*), and five genera showed a reduction in abundance (*Ralstonia, Pelomonas, Brevundimonas, Bosea*, and *Companilactobacillus*). These genera have been sporadically detected at low abundance in other *Drosophila* microbiome studies. Interestingly, *A. aceti* has been shown to be equally virulent to wild-type and immune-deficient flies, unlike other *Acetobacter* strains that were constrained by a specific host immune effector response [92]. While a small difference, the unique decrease in *A. aceti* presence, but not other *Acetobacter* strains, further suggests this species has unique host-microbe interaction properties relative to other *Acetobacter* species. Both genera whose presence was associated with a decrease in DCV viral load (*Bacillus, Streptococcus*) have appeared in past experimental studies in *Drosophila melanogaster*. The *Bacillus* species *B. thuringiensis* – a soil dwelling bacterium widely used as a microbial insecticide [93] – can cause highly virulent infections in *D. melanogaster* and is able to evade antimicrobial peptides (AMPs) [94]. *Streptococcus pneumoniae* and *S. agalactiae*, while not natural insect pathogens, induce strong Toll-dependent immune responses upon systemic infection in *D. melanogaster* and have been shown to prime the fly immune system through stimulation of haemocytes and Toll [95,96]. *Streptococcus* virulence in *D. melanogaster* depends on bacterial binding to host glycosaminoglycans, which downregulates *Drosophila* AMP production [96]. One potential mechanism mediating improved host defence against DCV could be microbe-mediated production of cyclic dinucleotides [97], that can prime host antiviral immunity to differing extents across *Drosophila* species [98]. However, the basis of this potential interaction between *Bacillus, Streptococcus*, and DCV, remains unclear.

Despite variation in both microbiome community composition and the presences and abundances of individual genera across our experimental data, this variation was only weakly explained by host phylogeny and host species identity. Instead, much of the variation occurred between replicates within species, suggesting that community composition in laboratory-reared *Drosophilidae* may be shaped predominantly by stochastic within-species effects. This is consistent with the transient and environmentally acquired nature of the *Drosophila* microbiome, where bacterial taxa are continually gained and lost through exposure to food, housing, and rearing conditions, and are weakly influenced by host genotype [25–27]. Correspondingly, we detected no significant interspecific correlations between DCV viral load and any alpha diversity metric, beta diversity axis, or individual bacterial taxa. Thus, in this study, bacterial community presence and relative relationships appear to be robust in the face of host virus infection. However, we observed a small number of significant correlations, suggesting that some variation in the microbiome is associated with RNA virus susceptibility, primarily through within-species or residual effects. While not included here, our methodology leaves the possibility open that viral infection induces shifts in total microbial load, rather than relative community structure [99]. Compared with other laboratory studies [22,25,29], our fly species harboured a broader set of bacterial genera and were not dominated primarily by *Acetobacter* and *Lactobacillaceae*. These differences may be due to differing geographic origins of our stocks, or a consequence of our husbandry conditions, including the use of multiple rearing diets and the parallel maintenance of a large number of species, which may have increased the diversity of incidental bacterial exposures.

Together, we find that microbiome composition in our panel of laboratory-reared *Drosophilidae* is weakly structured by host phylogeny, but that variation in specific bacterial taxa can be associated with meaningful differences in RNA virus replication across hosts. It remains unclear whether similar associations occur in wild *Drosophilidae*. Future experimental manipulation of the microbiome may potentially reveal causal roles for these taxa in shaping viral susceptibility.

