## Supplementary Text for "Examining interactions between the microbiome and viral infection across *Drosophila* species"

**Supplementary Materials**

The following document contains supplementary methods, tables, and figures for Imrie et al., Examining interactions between the microbiome and viral infection across *Drosophila* species (2026).

**Supplementary Tables**

| **Species** | **Genus** | **Diet** |
| --- | --- | --- |
| *D. affinis* | *Drosophila* | Malt |
| *D. americana* | *Drosophila* | Malt |
| *D. ananassae* | *Drosophila* | Cornmeal |
| *D. arizonae* | *Drosophila* | Banana |
| *D. baimaii* | *Drosophila* | Cornmeal |
| *D. buzzatii* | *Drosophila* | Malt |
| *D. euronotus* | *Drosophila* | Cornmeal |
| *D. flavomontana* | *Drosophila* | Malt w. additional yeast |
| *D. hydei* | *Drosophila* | Cornmeal |
| *D. immigrans* | *Drosophila* | Malt w. additional yeast |
| *D. lacicola* | *Drosophila* | Malt |
| *D. melanogaster* | *Drosophila* | Cornmeal |
| *D. montana* | *Drosophila* | Malt w. additional yeast |
| *D. nasuta* | *Drosophila* | Cornmeal |
| *D. nebulosa* | *Drosophila* | Cornmeal |
| *D. paramelanica* | *Drosophila* | Cornmeal |
| *D. persimilis* | *Drosophila* | Malt |
| *D. prosaltans* | *Drosophila* | Propionic |
| *D. pseudoobscura* | *Drosophila* | Malt |
| *D. putrida* | *Drosophila* | Propionic |
| *D. saltans* | *Drosophila* | Propionic |
| *D. santomea* | *Drosophila* | Cornmeal |
| *D. simulans* | *Drosophila* | Cornmeal |
| *D. sturtevanti* | *Drosophila* | Cornmeal |
| *D. subobscura* | *Drosophila* | Cornmeal |
| *D. sucinea* | *Drosophila* | Cornmeal |
| *D. teissieri* | *Drosophila* | Cornmeal |
| *D. virilis* | *Drosophila* | Propionic |
| *D. yakuba* | *Drosophila* | Cornmeal |
| *S. lebanonensis* | *Scaptodrosophila* | Propionic |
| *S. pattersoni* | *Scaptodrosophila* | Banana |
| *Z. tuberculatus* | *Zaprionous* | Banana |

**Supplementary Table 1: *Drosophilidae* host species and rearing diets.** Full recipes can be found at doi:10.6084/m9.figshare.21590724.v1. Some species were provided with additional dried yeast added to the top of their media once set.

|  | Uninjected | | | Saline Injected | | | DCV Infected | | |
| --- | --- | --- | --- | --- | --- | --- | --- | --- | --- |
| *D. affinis* | 12 | 10 | 12 | 11 | 10 | 10 | 10 | 11 | 10 |
| *D. americana* | 12 | 12 | 12 | 11 | 12 | 11 | 11 | 10 | 11 |
| *D. ananassae* | 12 | 11 | 12 | 11 | 12 | 11 | 11 | 12 | 11 |
| *D. arizonae* | 12 | 12 | 10 | 12 | 11 | 12 |  | 12 | 12 |
| *D. baimaii* | 12 | 12 | 12 | 12 | 10 | 9 | 10 | 11 | 11 |
| *D. buzzatii* | 10 | 11 | 9 | 10 | 11 | 9 | 12 | 11 | 9 |
| *D. euronotus* | 12 | 10 | 12 | 12 | 12 | 9 | 11 | 11 | 11 |
| *D. flavomontana* | 11 | 12 | 12 | 12 | 12 | 9 | 11 | 12 | 11 |
| *D. hydei* | 12 | 12 | 12 | 12 |  | 12 | 12 | 12 | 9 |
| *D. immigrans* | 12 | 11 | 11 | 10 | 12 | 11 | 12 | 11 | 12 |
| *D. lacicola* | 10 | 12 | 11 | 12 | 11 | 11 | 11 | 11 | 11 |
| *D. melanogaster* | 12 | 12 | 12 |  | 12 | 11 | 11 | 12 | 12 |
| *D. montana* | 12 | 12 | 12 | 12 | 11 | 10 | 11 | 12 | 12 |
| *D. nasuta* | 11 | 12 | 11 |  | 11 | 9 | 11 | 12 | 11 |
| *D. nebulosa* |  | 12 | 12 | 11 | 10 | 8 | 10 | 11 | 10 |
| *D. paramelanica* | 11 | 12 | 11 | 11 | 11 | 8 | 11 | 12 | 12 |
| *D. persimilis* | 10 | 11 | 11 |  | 11 | 8 | 12 | 12 | 10 |
| *D. prosaltans* | 12 | 12 | 12 | 11 | 11 | 10 | 12 | 11 | 12 |
| *D. pseudoobscura* | 11 | 12 | 10 |  | 10 | 10 |  | 11 | 12 |
| *D. putrida* | 12 | 12 | 12 | 12 | 12 | 11 | 10 | 10 | 11 |
| *D. saltans* | 10 | 12 | 12 |  | 9 | 11 | 10 | 12 | 12 |
| *D. santomea* | 12 | 12 | 12 | 11 | 12 | 11 |  | 11 | 11 |
| *D. simulans* |  | 12 | 12 | 12 | 12 | 10 | 11 | 12 | 9 |
| *D. sturtevanti* | 12 | 12 | 9 | 11 | 11 | 10 |  | 10 | 9 |
| *D. subobscura* | 12 | 11 | 11 | 10 | 12 |  |  | 12 | 12 |
| *D. sucinea* |  | 12 | 12 | 12 | 12 | 12 |  | 11 | 12 |
| *D. teissieri* |  | 11 | 12 | 12 | 9 | 10 |  | 10 | 9 |
| *D. virilis* | 12 |  | 12 | 12 | 11 | 11 | 12 | 9 | 12 |
| *D. yakuba* | 12 | 11 | 10 | 10 | 12 | 8 | 11 | 11 | 8 |
| *S. lebanonensis* | 11 | 12 | 12 |  | 10 | 12 | 10 | 10 | 12 |
| *S. pattersoni* | 12 | 11 | 12 | 12 | 12 | 12 | 11 | 12 | 11 |
| *Z. tuberculatus* | 12 | 12 | 11 | 12 | 12 | 11 | 12 | 11 | 10 |

**Supplementary Table 2: Biological replicates (vials) and flies per vial by species and injection condition.** Black boxes indicate replicates that were excluded from analysis, all of which were removed due to low 16S read count (<1,350).

| **Direction** | **Name** | **Sequence** |
| --- | --- | --- |
| Forward | DCV_F | GACACTGCCTTTGATTAG |
|  | RPL32_F-a | TGCCAAGTTGTCGCACAAATGG |
|  | RPL32_F-b | TGCTAAGTTGTCGCACAAATGG |
|  | RPL32_F-c | TGCCAAGCTGTCGCACAAATGG |
|  | RPL32_F-d | TGCTAAGCTGTCGCACAAATGG |
|  | RPL32_F-e | TGCGAAGTTGTCGCACAAATGG |
|  | RPL32_F-f | TGCGAAGCTGTCGCACAAATGG |
| Reverse | DCV_R | CCCTCTGGGAACTAAATG |
|  | RPL32_R-a | TGCGCTTGTTGGAACCGTAAC |
|  | RPL32_R-b | TGCGCTTGTTGGATCCGTAAC |
|  | RPL32_R-c | TGCGCTTGTTGGAACCATAAC |
|  | RPL32_R-d | TGCGCTTGTTGGAGCCGTAAC |
|  | RPL32_R-e | TGCGCTTGTTAGAACCGTAAC |
|  | RPL32_R-f | TACGCTTGTTGGAACCGTAAC |
|  | RPL32_R-g | TGCGCTTGTTGGAACCGTAGC |
|  | RPL32_R-h | TGCGCTTGTTCGATCCGTAAC |
|  | RPL32_R-i | TGCGCTTGTTGGAGCCATAAC |
|  | RPL32_R-j | TGCGCTTGTTTGATCCGTAAC |
|  | RPL32_R-k | TGCGCTTGTTTGAACCATAAC |
|  | RPL32_R-l | TACGCTTGTTGGAACCATAAC |
|  | RPL32_R-m | TACGCTTGTTGGAGCCGTAAC |
|  | RPL32_R-n | TGCGCTGGTTGGAACCATAAC |
|  | RPL32_R-o | TGAGCTTGTTCGATCCGTAAC |
|  | RPL32_R-p | TACGCTTGTTGGAGCCATAAC |
|  | RPL32_R-q | TGAGCTTGTTTGATCCGTAAC |
|  | RPL32_R-r | TAAGCTTGTTGGATCCGTAGC |
|  | RPL32_R-s | TCAGCTTGTTGGATCCATAGC |

**Supplementary Table 3: DCV and RPL32 q-PCR primers**

| **Species** | **Forward** | **Reverse** |
| --- | --- | --- |
| *D. affinis* | F-a | R-i |
| *D. americana* | F-c | R-a |
| *D. ananassae* | F-f | R-a |
| *D. arizonae* | F-a | R-a |
| *D. baimaii* | F-a | R-r |
| *D. buzzati* | F-a | R-e |
| *D. erecta* | F-d | R-h |
| *D. euronotus* | F-a | R-g |
| *D. flavomontana* | F-c | R-a |
| *D. hydei* | F-a | R-a |
| *D. immigrans* | F-b | R-p |
| *D. lacicola* | F-c | R-a |
| *D. melanogaster* | F-d | R-h |
| *D. montana* | F-c | R-a |
| *D. nasuta* | F-b | R-f |
| *D. nebulosa* | F-b | R-c |
| *D. paramelanica* | F-a | R-g |
| *D. persimilis* | F-a | R-b |
| *D. prosaltans* | F-a | R-n |
| *D. pseudoobscura* | F-a | R-m |
| *D. putridia* | F-d | R-q |
| *D. saltans* | F-a | R-n |
| *D. santomea* | F-a | R-n |
| *D. simulans* | F-d | R-h |
| *D. sturtevanti* | F-a | R-l |
| *D. subobscura* | F-a | R-i |
| *D. sucinea* | F-b | R-k |
| *D. teissieri* | F-d | R-h |
| *D. virilis* | F-c | R-a |
| D. yakuba | F-d | R-h |
| S. lebanonensis | F-d | R-h |
| S. pattersoni | F-a | R-m |
| Z. tuberculatus | F-a | R-c |

**Supplementary Table 4: RPL32 Primer Combinations**

| **Cycle step** | **Temp.** | **Time/rate** | **Cycles** |
| --- | --- | --- | --- |
| Initial denaturation | 95˚C | 2 min | 1 |
| Denaturation | 95˚C | 5 sec | ) 40 |
| Annealing/Extension | 60˚C | 15 sec |  |
| Melt curve | 60˚C - 95˚C | 0.1˚C/s | 1 |

**Supplementary Table 5: q(RT)-PCR Cycle Conditions**

| **Response** | **Model Structure** | **Diet (Proportion of Total Variance)** |
| --- | --- | --- |
| Richness | 1 | 0.05 (0.00, 0.20) |
| Richness | 2 | 0.06 (0.00, 0.24) |
| Evenness | 1 | 0.09 (0.00, 0.31) |
| Evenness | 2 | 0.09 (0.00, 0.30) |
| Shannon Diversity | 1 | 0.07 (0.00, 0.27) |
| Shannon Diversity | 2 | 0.07 (0.00, 0.26) |
| PC1 | 1 | 0.01 (0.00, 0.03) |
| PC1 | 2 | 0.01 (0.00, 0.03) |
| PC2 | 1 | 0.21 (0.01, 0.59) |
| PC2 | 2 | 0.21 (0.00, 0.58) |
| PC3 | 1 | 0.05 (0.00, 0.22) |
| PC3 | 2 | 0.05 (0.00, 0.21) |
| PC4 | 1 | 0.01 (0.00, 0.04) |
| PC4 | 2 | 0.01 (0.00, 0.03) |
| NMDS1 | 1 | 0.01 (0.00, 0.02) |
| NMDS1 | 2 | 0.01 (0.00, 0.03) |
| NMDS2 | 1 | 0.17 (0.01, 0.51) |
| NMDS2 | 2 | 0.18 (0.01, 0.50) |
| NMDS3 | 1 | 0.07 (0.00, 0.29) |
| NMDS3 | 2 | 0.09 (0.00, 0.31) |
| NMDS4 | 1 | 0.01 (0.00, 0.04) |
| NMDS4 | 2 | 0.01 (0.00, 0.04) |
| Presence | 1 | 0.02 (0.00, 0.10) |
| Presence | 2 | 0.02 (0.00, 0.08) |
| Abundance | 1 | 0.00 (0.00, 0.02) |
| Abundance | 2 | 0.00 (0.00, 0.02) |

**Supplementary Table 6: Estimates of the proportion of total variance in different response variables explained by host rearing diet. Proportions whose 95% CIs do not overlap 0.05 are highlighted in bold.**

| **#** | **Taxa** | **Phylogeny** | **Species-specific** | **Residuals** |
| --- | --- | --- | --- | --- |
| 01 | *Serratia* | 0.08 (0.00, 0.26) | 0.11 (0.00, 0.30) | **0.81 (0.60, 1.00)** |
| 02 | *Serratia* | 0.33 (0.00, 0.80) | 0.13 (0.00, 0.48) | **0.54 (0.13, 0.98)** |
| 03 | *Escherichia-Shigella* | 0.03 (0.00, 0.10) | 0.03 (0.00, 0.10) | **0.95 (0.84, 1.00)** |
| 04 | *Buttiauxella* | 0.05 (0.00, 0.19) | 0.05 (0.00, 0.16) | **0.90 (0.75, 1.00)** |
| 05 | *Buttiauxella* | 0.18 (0.00, 0.45) | 0.07 (0.00, 0.25) | **0.74 (0.49, 0.98)** |
| 06 | *Buttiauxella* | 0.10 (0.00, 0.35) | 0.14 (0.00, 0.35) | **0.76 (0.52, 1.00)** |
| 07 | *Providencia rettgeri* | 0.20 (0.00, 0.45) | 0.15 (0.00, 0.38) | **0.66 (0.42, 0.87)** |
| 08 | *Providencia* | 0.24 (0.00, 0.52) | 0.12 (0.00, 0.37) | **0.64 (0.39, 0.88)** |
| 09 | *Providencia* | 0.04 (0.00, 0.14) | 0.03 (0.00, 0.10) | **0.93 (0.82, 1.00)** |
| 10 | *Providencia* | 0.20 (0.00, 0.47) | 0.10 (0.00, 0.32) | **0.70 (0.43, 0.93)** |
| 11 | *Providencia* | 0.23 (0.00, 0.53) | 0.12 (0.00, 0.36) | **0.65 (0.38, 0.91)** |
| 12 | *Providencia* | 0.11 (0.00, 0.31) | 0.05 (0.00, 0.18) | **0.84 (0.64, 1.00)** |
| 13 | *Enhydrobacter aerosaccus* | 0.14 (0.00, 0.44) | 0.10 (0.00, 0.34) | **0.75 (0.46, 1.00)** |
| 14 | *Pseudomonas* | 0.14 (0.00, 0.32) | 0.05 (0.00, 0.17) | **0.82 (0.62, 0.99)** |
| 15 | *Stenotrophomonas maltophilia* | 0.09 (0.00, 0.28) | 0.07 (0.00, 0.23) | **0.84 (0.63, 1.00)** |
| 16 | *Stenotrophomonas maltophilia* | 0.26 (0.00, 0.76) | 0.39 (0.00, 0.76) | **0.35 (0.07, 0.64)** |
| 17 | *Pelomonas* | 0.07 (0.00, 0.22) | 0.06 (0.00, 0.18) | **0.88 (0.72, 1.00)** |
| 18 | *Ralstonia* | 0.10 (0.00, 0.30) | 0.07 (0.00, 0.21) | **0.83 (0.64, 0.98)** |
| 19 | *Acetobacter aceti* | 0.12 (0.00, 0.38) | 0.10 (0.00, 0.27) | **0.78 (0.56, 0.99)** |
| 20 | *Acetobacter sicerae* | 0.29 (0.00, 0.70) | 0.36 (0.00, 0.69) | **0.35 (0.12, 0.60)** |
| 21 | *Acetobacter* | 0.06 (0.00, 0.21) | 0.11 (0.00, 0.30) | **0.83 (0.61, 1.00)** |
| 22 | *Acetobacter indonesiensis* | 0.06 (0.00, 0.22) | 0.04 (0.00, 0.14) | **0.90 (0.74, 1.00)** |
| 23 | *Acetobacter* | 0.09 (0.00, 0.28) | 0.04 (0.00, 0.15) | **0.86 (0.68, 1.00)** |
| 24 | *Commensalibacter* | 0.13 (0.00, 0.41) | 0.14 (0.00, 0.39) | **0.74 (0.45, 1.00)** |
| 25 | *Commensalibacter* | 0.06 (0.00, 0.22) | 0.05 (0.00, 0.19) | **0.89 (0.68, 1.00)** |
| 26 | *Gluconobacter* | 0.05 (0.00, 0.20) | 0.04 (0.00, 0.16) | **0.90 (0.72, 1.00)** |
| 27 | *Brucella* | 0.09 (0.00, 0.33) | 0.11 (0.00, 0.33) | **0.80 (0.54, 1.00)** |
| 28 | *Brevundimonas* | 0.09 (0.00, 0.23) | 0.04 (0.00, 0.14) | **0.87 (0.72, 1.00)** |
| 29 | *Afipia* | 0.21 (0.00, 0.58) | 0.19 (0.00, 0.53) | **0.60 (0.24, 0.94)** |
| 30 | *Bosea* | 0.06 (0.00, 0.18) | 0.04 (0.00, 0.13) | **0.90 (0.77, 1.00)** |
| 31 | *Bacteroides* | 0.15 (0.00, 0.48) | 0.24 (0.00, 0.55) | **0.62 (0.29, 0.95)** |
| 32 | *Dysgonomonas* | 0.05 (0.00, 0.19) | 0.06 (0.00, 0.19) | **0.90 (0.72, 1.00)** |
| 33 | *Sediminibacterium salmoneum* | 0.04 (0.00, 0.14) | 0.04 (0.00, 0.15) | **0.92 (0.78, 1.00)** |
| 34 | *Levilactobacillus* | 0.19 (0.00, 0.43) | 0.07 (0.00, 0.23) | **0.74 (0.51, 0.94)** |
| 35 | *Levilactobacillus* | 0.14 (0.00, 0.50) | 0.29 (0.00, 0.61) | **0.57 (0.21, 0.90)** |
| 36 | *Fructilactobacillus* | 0.35 (0.00, 0.70) | 0.21 (0.00, 0.53) | **0.44 (0.17, 0.72)** |
| 37 | *Companilactobacillus* | 0.08 (0.00, 0.29) | 0.36 (0.00, 0.65) | **0.56 (0.24, 0.89)** |
| 38 | *Lactobacillus* | 0.04 (0.00, 0.14) | 0.06 (0.00, 0.20) | **0.90 (0.74, 1.00)** |
| 39 | *Staphylococcus* | 0.07 (0.00, 0.22) | 0.04 (0.00, 0.15) | **0.89 (0.73, 1.00)** |
| 40 | *Bacillus* | 0.05 (0.00, 0.18) | 0.05 (0.00, 0.17) | **0.90 (0.75, 1.00)** |
| 41 | *Enterococcus faecalis* | 0.08 (0.00, 0.27) | 0.08 (0.00, 0.26) | **0.84 (0.62, 1.00)** |
| 42 | *Streptococcus* | 0.28 (0.00, 0.62) | 0.10 (0.00, 0.35) | **0.62 (0.27, 0.93)** |
| 43 | *Streptococcus* | 0.11 (0.00, 0.35) | 0.14 (0.00, 0.36) | **0.75 (0.50, 0.98)** |
| 44 | *Finegoldia magna* | 0.16 (0.00, 0.51) | 0.18 (0.00, 0.47) | **0.67 (0.35, 0.98)** |
| 45 | *Corynebacterium* | 0.29 (0.00, 0.61) | 0.11 (0.00, 0.38) | **0.61 (0.28, 0.92)** |
| 46 | *Corynebacterium nuruki* | 0.07 (0.00, 0.25) | 0.09 (0.00, 0.23) | **0.84 (0.66, 1.00)** |
| 47 | *Corynebacterium* | 0.03 (0.00, 0.13) | 0.03 (0.00, 0.10) | **0.94 (0.82, 1.00)** |
| 48 | *Lawsonella* | 0.10 (0.00, 0.31) | 0.09 (0.00, 0.27) | **0.81 (0.60, 1.00)** |
| 49 | *Rhodococcus* | 0.13 (0.00, 0.36) | 0.07 (0.00, 0.26) | **0.80 (0.55, 1.00)** |
| 50 | *Cutibacterium* | 0.10 (0.00, 0.25) | 0.05 (0.00, 0.17) | **0.85 (0.70, 1.00)** |

**Supplementary Table 7: Estimates of the proportion of total variance in bacterial ASV presence explained by host phylogeny, a species-specific categorical effect, and the model residuals. Proportions whose 95% CIs do not overlap 0.05 are highlighted in bold.**

| **#** | **Taxa** | **Phylogeny** | **Species-specific** | **Residuals** |
| --- | --- | --- | --- | --- |
| 01 | *Serratia* | 0.07 (0.00, 0.21) | 0.06 (0.00, 0.17) | **0.87 (0.74, 1.00)** |
| 02 | *Serratia* | 0.10 (0.00, 0.29) | 0.10 (0.00, 0.23) | **0.80 (0.65, 0.94)** |
| 03 | *Escherichia-Shigella* | 0.08 (0.00, 0.28) | 0.09 (0.00, 0.30) | **0.84 (0.57, 1.00)** |
| 04 | *Buttiauxella* | 0.07 (0.00, 0.24) | 0.07 (0.00, 0.20) | **0.85 (0.69, 1.00)** |
| 05 | *Buttiauxella* | 0.11 (0.00, 0.35) | 0.09 (0.00, 0.29) | **0.81 (0.54, 1.00)** |
| 06 | *Buttiauxella* | 0.25 (0.00, 0.66) | 0.21 (0.00, 0.58) | **0.54 (0.19, 0.94)** |
| 07 | *Providencia rettgeri* | 0.14 (0.00, 0.43) | 0.19 (0.00, 0.49) | **0.67 (0.37, 0.95)** |
| 08 | *Providencia* | 0.09 (0.00, 0.34) | 0.11 (0.00, 0.36) | **0.79 (0.51, 1.00)** |
| 09 | *Providencia* | 0.23 (0.00, 0.46) | 0.07 (0.00, 0.25) | **0.70 (0.48, 0.91)** |
| 10 | *Providencia* | 0.17 (0.00, 0.52) | 0.17 (0.00, 0.48) | **0.66 (0.35, 0.99)** |
| 11 | *Providencia* | 0.21 (0.00, 0.59) | 0.18 (0.00, 0.49) | **0.61 (0.30, 0.97)** |
| 12 | *Providencia* | 0.25 (0.00, 0.74) | 0.28 (0.00, 0.68) | **0.47 (0.13, 0.91)** |
| 13 | *Enhydrobacter aerosaccus* | 0.43 (0.00, 0.91) | 0.24 (0.00, 0.76) | **0.33 (0.01, 0.88)** |
| 14 | *Pseudomonas* | 0.07 (0.00, 0.18) | 0.03 (0.00, 0.12) | **0.90 (0.78, 1.00)** |
| 15 | *Stenotrophomonas maltophilia* | 0.16 (0.00, 0.50) | 0.22 (0.00, 0.54) | **0.62 (0.31, 0.98)** |
| 16 | *Stenotrophomonas maltophilia* | 0.28 (0.00, 0.69) | 0.17 (0.00, 0.56) | **0.55 (0.18, 0.97)** |
| 17 | *Pelomonas* | 0.10 (0.00, 0.29) | 0.06 (0.00, 0.21) | **0.84 (0.65, 1.00)** |
| 18 | *Ralstonia* | 0.15 (0.00, 0.34) | 0.04 (0.00, 0.16) | **0.81 (0.63, 0.99)** |
| 19 | *Acetobacter aceti* | 0.16 (0.00, 0.42) | 0.16 (0.00, 0.35) | **0.68 (0.49, 0.86)** |
| 20 | *Acetobacter sicerae* | 0.14 (0.00, 0.50) | 0.16 (0.00, 0.48) | **0.70 (0.34, 1.00)** |
| 21 | *Acetobacter* | 0.06 (0.00, 0.21) | 0.16 (0.00, 0.30) | **0.78 (0.61, 0.93)** |
| 22 | *Acetobacter indonesiensis* | 0.07 (0.00, 0.24) | 0.05 (0.00, 0.17) | **0.87 (0.71, 1.00)** |
| 23 | *Acetobacter* | 0.07 (0.00, 0.23) | 0.06 (0.00, 0.20) | **0.87 (0.70, 1.00)** |
| 24 | *Commensalibacter* | 0.15 (0.00, 0.37) | 0.12 (0.00, 0.29) | **0.74 (0.56, 0.91)** |
| 25 | *Commensalibacter* | **0.28 (0.08, 0.50)** | 0.04 (0.00, 0.15) | **0.68 (0.50, 0.85)** |
| 26 | *Gluconobacter* | 0.21 (0.00, 0.60) | 0.14 (0.00, 0.48) | **0.65 (0.29, 1.00)** |
| 27 | *Brucella* | 0.16 (0.00, 0.52) | 0.15 (0.00, 0.49) | **0.69 (0.33, 1.00)** |
| 28 | *Brevundimonas* | 0.07 (0.00, 0.22) | 0.05 (0.00, 0.16) | **0.88 (0.73, 1.00)** |
| 29 | *Afipia* | 0.26 (0.00, 0.75) | 0.23 (0.00, 0.67) | **0.51 (0.13, 1.00)** |
| 30 | *Bosea* | 0.08 (0.00, 0.25) | 0.05 (0.00, 0.17) | **0.87 (0.69, 1.00)** |
| 31 | *Bacteroides* | 0.14 (0.00, 0.47) | 0.15 (0.00, 0.47) | **0.71 (0.36, 1.00)** |
| 32 | *Dysgonomonas* | 0.08 (0.00, 0.25) | 0.05 (0.00, 0.16) | **0.87 (0.72, 1.00)** |
| 33 | *Sediminibacterium salmoneum* | 0.16 (0.00, 0.54) | 0.17 (0.00, 0.52) | **0.67 (0.31, 1.00)** |
| 34 | *Levilactobacillus* | 0.22 (0.00, 0.49) | 0.13 (0.00, 0.35) | **0.65 (0.42, 0.86)** |
| 35 | *Levilactobacillus* | 0.18 (0.00, 0.54) | 0.13 (0.00, 0.43) | **0.70 (0.34, 1.00)** |
| 36 | *Fructilactobacillus* | 0.25 (0.00, 0.66) | 0.25 (0.00, 0.64) | **0.51 (0.16, 0.91)** |
| 37 | *Companilactobacillus* | 0.27 (0.00, 0.66) | 0.18 (0.00, 0.55) | **0.56 (0.20, 0.95)** |
| 38 | *Lactobacillus* | 0.10 (0.00, 0.35) | 0.15 (0.00, 0.39) | **0.76 (0.49, 1.00)** |
| 39 | *Staphylococcus* | 0.24 (0.00, 0.60) | 0.19 (0.00, 0.55) | **0.57 (0.24, 0.95)** |
| 40 | *Bacillus* | 0.03 (0.00, 0.11) | 0.03 (0.00, 0.12) | **0.94 (0.82, 1.00)** |
| 41 | *Enterococcus faecalis* | 0.20 (0.00, 0.63) | 0.15 (0.00, 0.49) | **0.65 (0.27, 1.00)** |
| 42 | *Streptococcus* | 0.18 (0.00, 0.59) | 0.21 (0.00, 0.60) | **0.61 (0.23, 1.00)** |
| 43 | *Streptococcus* | 0.35 (0.00, 0.69) | 0.17 (0.00, 0.54) | **0.49 (0.17, 0.82)** |
| 44 | *Finegoldia magna* | 0.21 (0.00, 0.63) | 0.14 (0.00, 0.48) | **0.65 (0.26, 1.00)** |
| 45 | *Corynebacterium* | 0.20 (0.00, 0.60) | 0.13 (0.00, 0.45) | **0.67 (0.29, 1.00)** |
| 46 | *Corynebacterium nuruki* | 0.12 (0.00, 0.33) | 0.04 (0.00, 0.16) | **0.83 (0.63, 1.00)** |
| 47 | *Corynebacterium* | 0.09 (0.00, 0.35) | 0.15 (0.00, 0.46) | **0.76 (0.43, 1.00)** |
| 48 | *Lawsonella* | 0.06 (0.00, 0.23) | 0.05 (0.00, 0.19) | **0.89 (0.69, 1.00)** |
| 49 | *Rhodococcus* | 0.27 (0.00, 0.74) | 0.25 (0.00, 0.70) | **0.48 (0.10, 0.97)** |
| 50 | *Cutibacterium* | 0.08 (0.00, 0.25) | 0.09 (0.00, 0.25) | **0.83 (0.64, 1.00)** |

**Supplementary Table 8: Estimates of the proportion of total variance in bacterial ASV CLR-transformed abundance explained by host phylogeny, a species-specific categorical effect, and the model residuals. Proportions whose 95% CIs do not overlap 0.05 are highlighted in bold.**

|  | **Before Infection (Uninjected)** | | **During Infection** | |
| --- | --- | --- | --- | --- |
| **Metric/Axis** | **R (phylogenetic)** | **R (marginal)** | **R (phylogenetic)** | **R (marginal)** |
| Richness | -0.10 (-0.93, 0.77) | -0.11 (-0.36, 0.12) | -0.14 (-0.94, 0.77) | -0.11 (-0.36, 0.14) |
| Evenness | -0.09 (-0.90, 0.78) | -0.03 (-0.28, 0.23) | -0.02 (-0.84, 0.81) | -0.05 (-0.34, 0.25) |
| Shannon Diversity | -0.10 (-0.94, 0.76) | -0.07 (-0.33, 0.21) | -0.05 (-0.86, 0.80) | -0.06 (-0.34, 0.22) |
| PC1 | -0.06 (-0.93, 0.80) | -0.16 (-0.40, 0.07) | -0.12 (-0.97, 0.71) | -0.05 (-0.28, 0.21) |
| PC2 | 0.07 (-0.75, 0.89) | 0.08 (-0.22, 0.36) | 0.21 (-0.62, 0.95) | 0.06 (-0.26, 0.43) |
| PC3 | 0.13 (-0.70, 0.87) | 0.01 (-0.36, 0.39) | -0.03 (-0.91, 0.82) | 0.03 (-0.19, 0.24) |
| PC4 | -0.09 (-0.93, 0.78) | -0.18 (-0.44, 0.06) | -0.04 (-0.86, 0.86) | -0.08 (-0.41, 0.26) |
| NMDS1 | -0.02 (-0.89, 0.86) | -0.14 (-0.36, 0.09) | -0.08 (-0.95, 0.78) | -0.05 (-0.27, 0.19) |
| NMDS2 | -0.22 (-0.99, 0.64) | -0.13 (-0.41, 0.12) | -0.38 (-0.99, 0.43) | -0.11 (-0.44, 0.25) |
| NMDS3 | -0.15 (-0.92, 0.68) | -0.03 (-0.37, 0.32) | 0.03 (-0.83, 0.91) | -0.04 (-0.26, 0.16) |
| NMDS4 | -0.22 (-0.97, 0.67) | -0.22 (-0.46, 0.01) | 0.05 (-0.78, 0.91) | -0.03 (-0.32, 0.26) |

**Supplementary Table 9: Interspecific and marginal correlations between DCV viral load and alpha diversity metrics, principle component (PC) or non-metric multidimensional scaling (NMDS) axes. Posteriors whose 95% CIs do not overlap 0 are highlighted in bold.**

**Supplementary Figures**

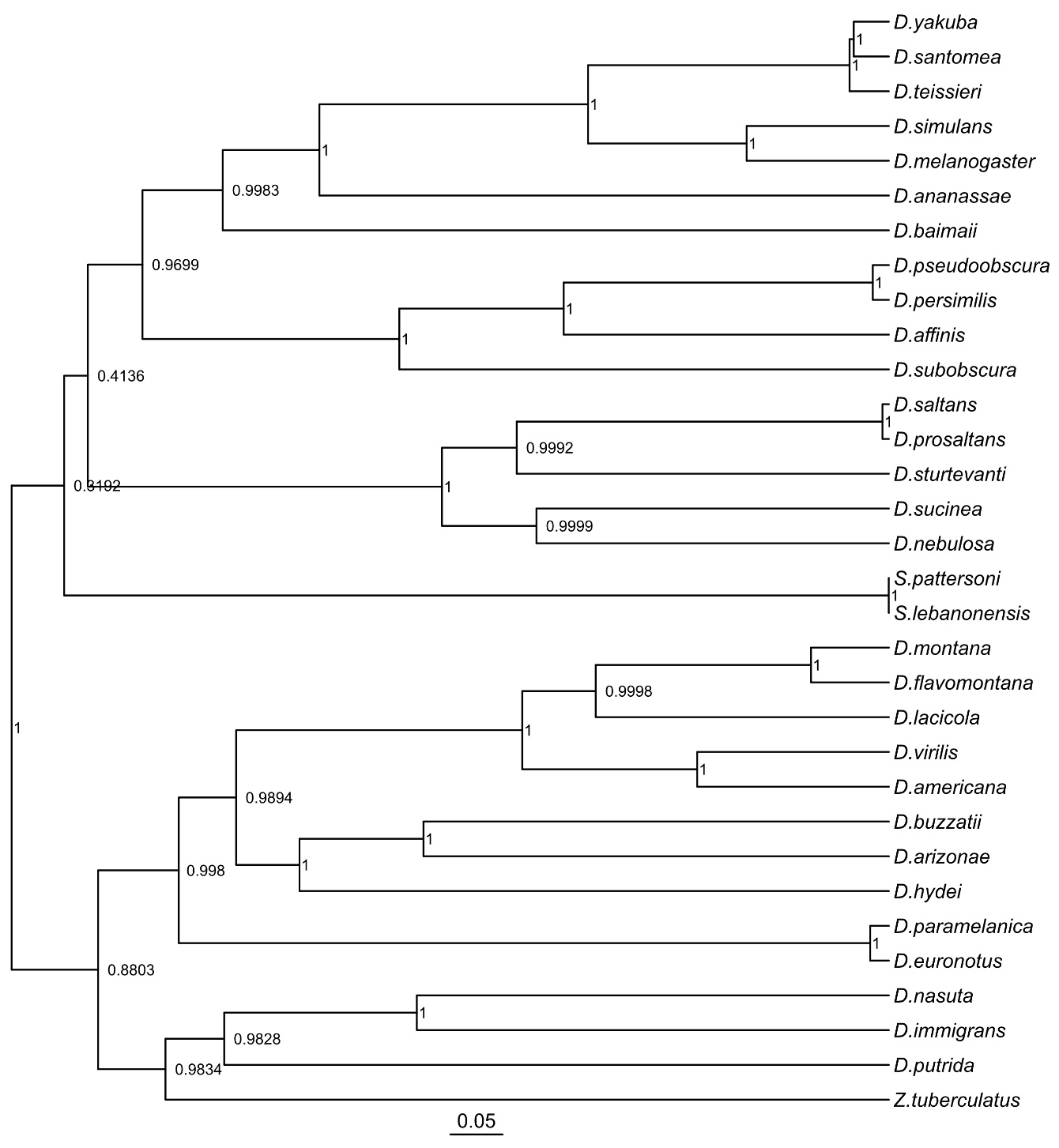

**Supplementary Figure 1: Phylogeny of *Drosophilidae* host species.** Evolutionary relationships are presented as a midpoint rooted, maximum clade credibility tree. Node labels are the posterior probabilities of each clade, and the scale bar represents nucleotide substitutions per site.

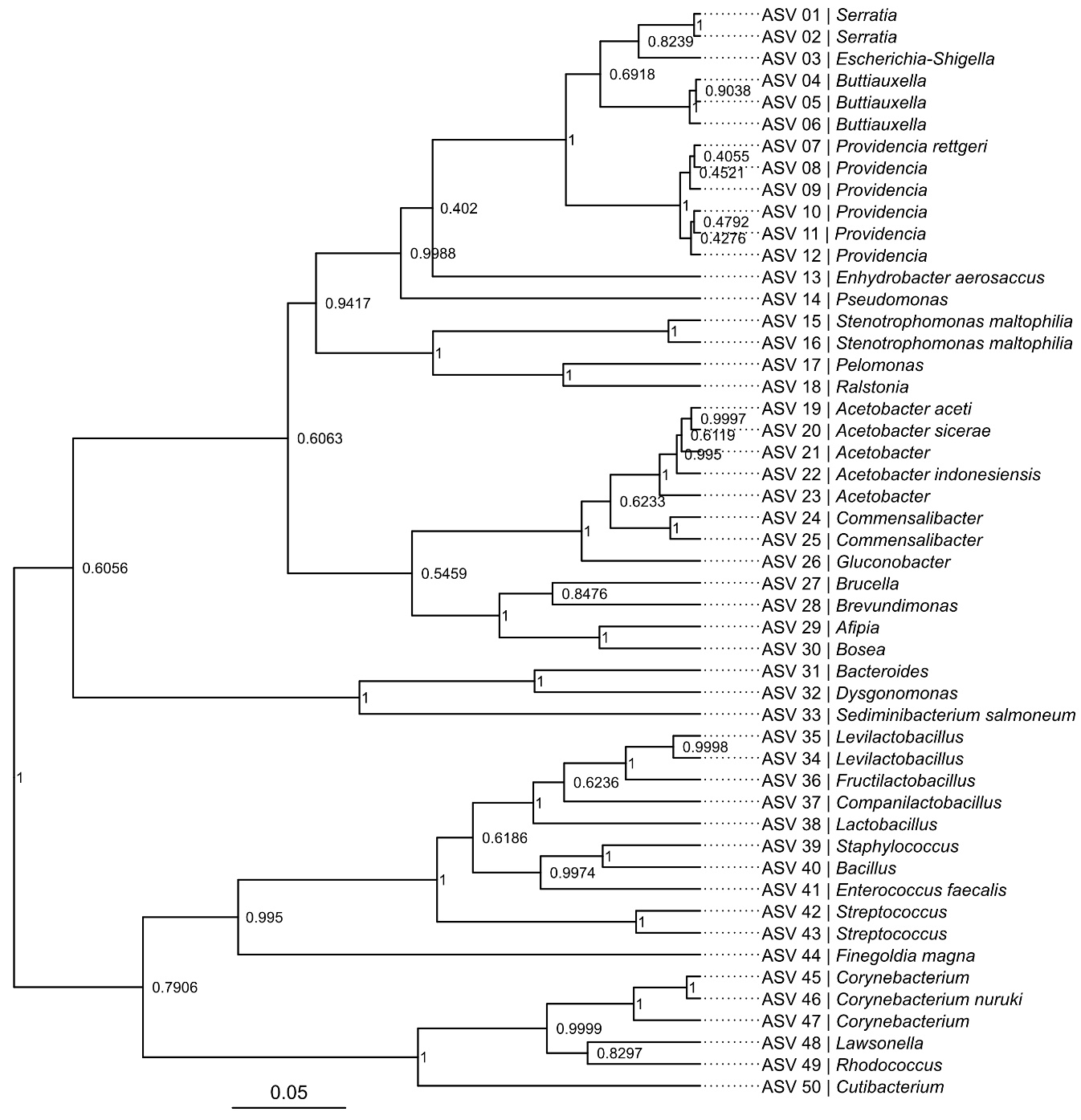

**Supplementary Figure 2: Phylogeny of bacterial ASVs.** Evolutionary relationships are presented as a midpoint rooted, maximum clade credibility tree. Node labels are the posterior probabilities of each clade, and the scale bar represents nucleotide substitutions per site.

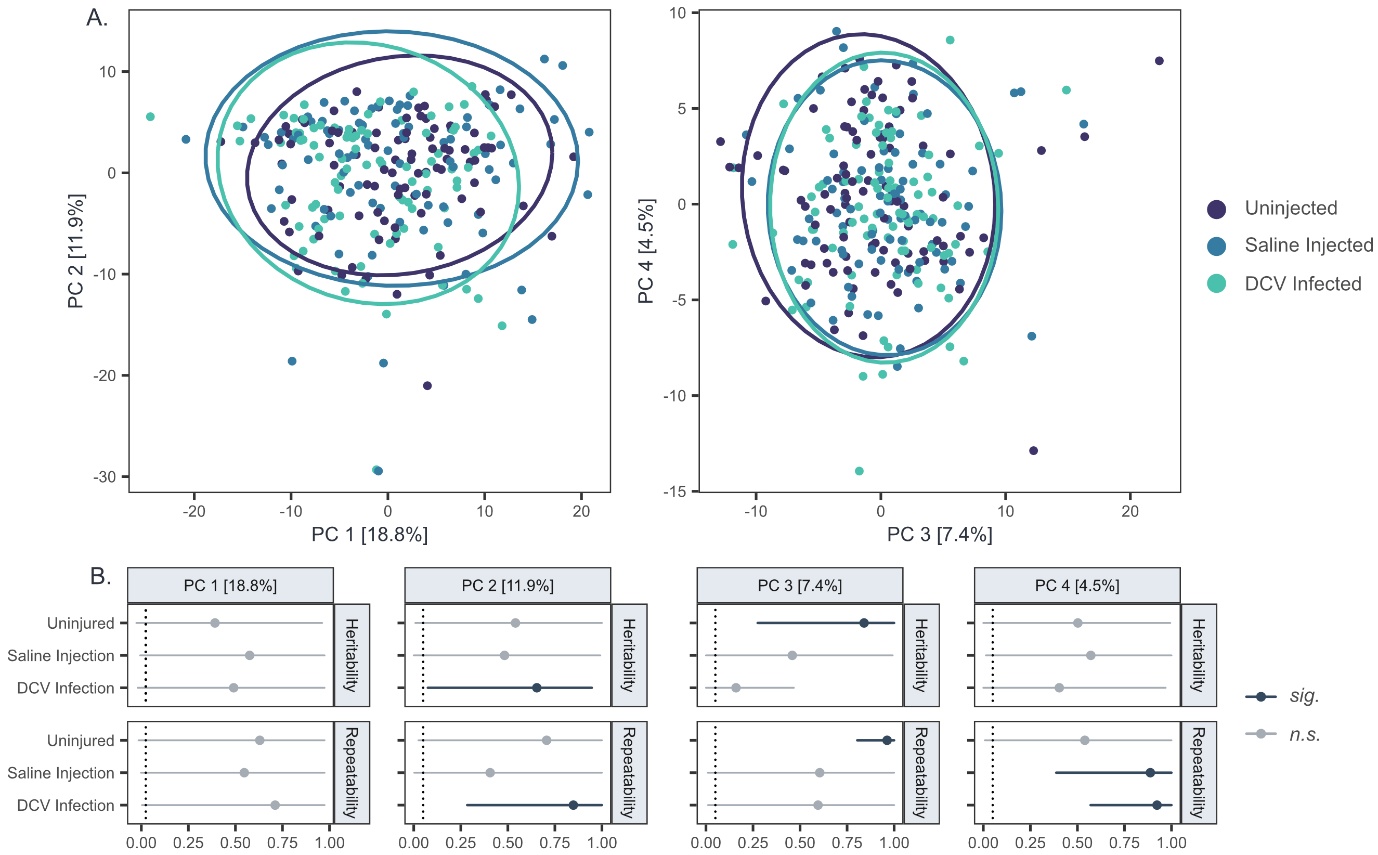

**Supplementary Figure 3: Principle Component Analysis (PCA) of bacterial ASV abundances across host species.** A) Scatter plots and ellipses for PC1 x PC2 (left) and PC3 x PC4 (right) by experimental condition. B) Estimates of phylogenetic heritability (top facet row) and repeatability (bottom facet row) for each PC axis under each experimental condition. Points represent the mean and bars the 95% CIs of the posterior distributions of each estimate. Non-zero estimates are highlighted in black.

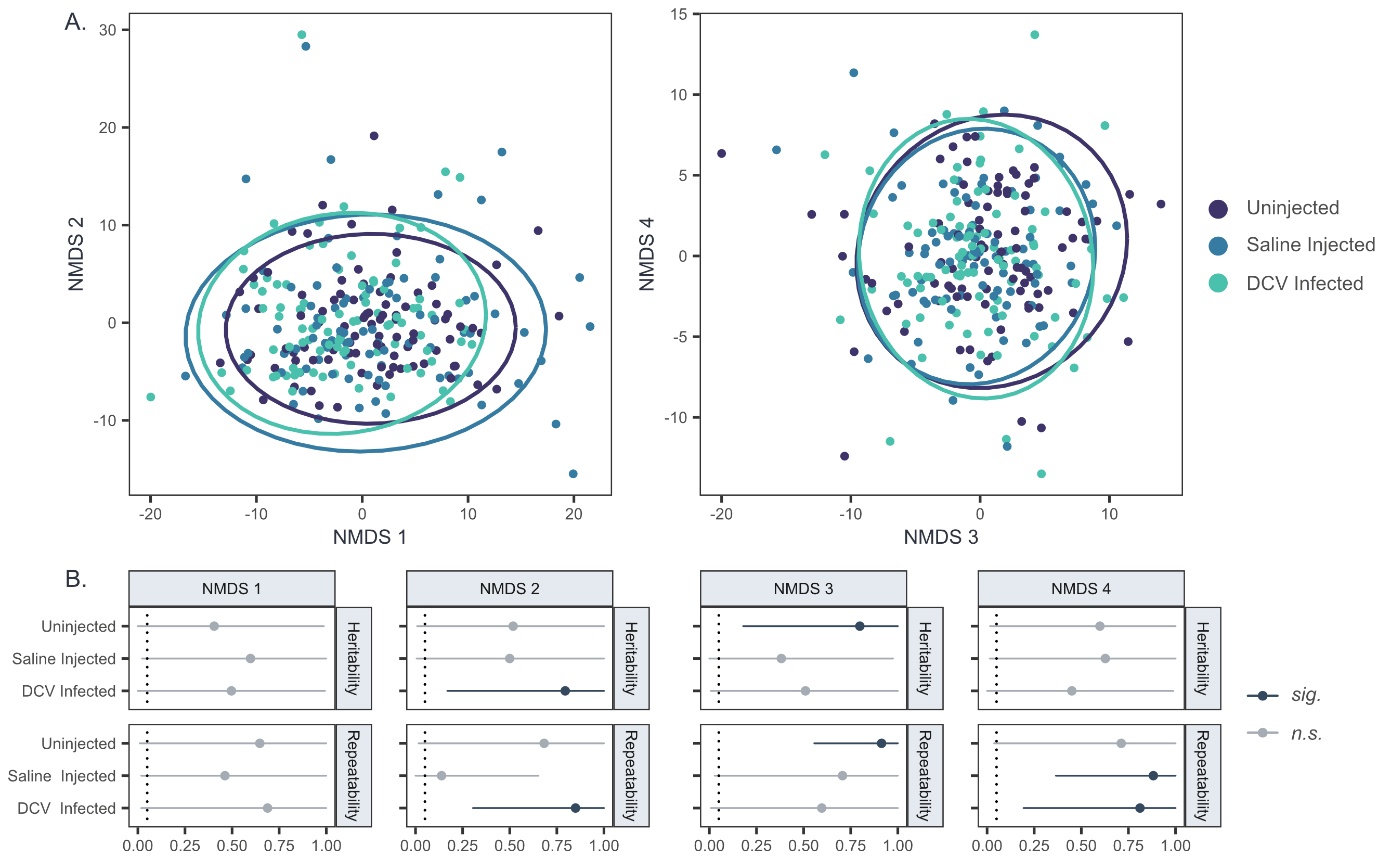

**Supplementary Figure 4: Non-Metric Multidimensional Scaling (NMDS) of bacterial ASV abundances across host species.** A) Scatter plots and ellipses for NMDS axes 1 x 2 (left) and 3 x 4 (right) by experimental condition. B) Estimates of phylogenetic heritability (top facet row) and repeatability (bottom facet row) for each NMDS axis under each experimental condition. Points represent the mean and bars the 95% CIs of the posterior distributions of each estimate. Non-zero estimates are highlighted in black.

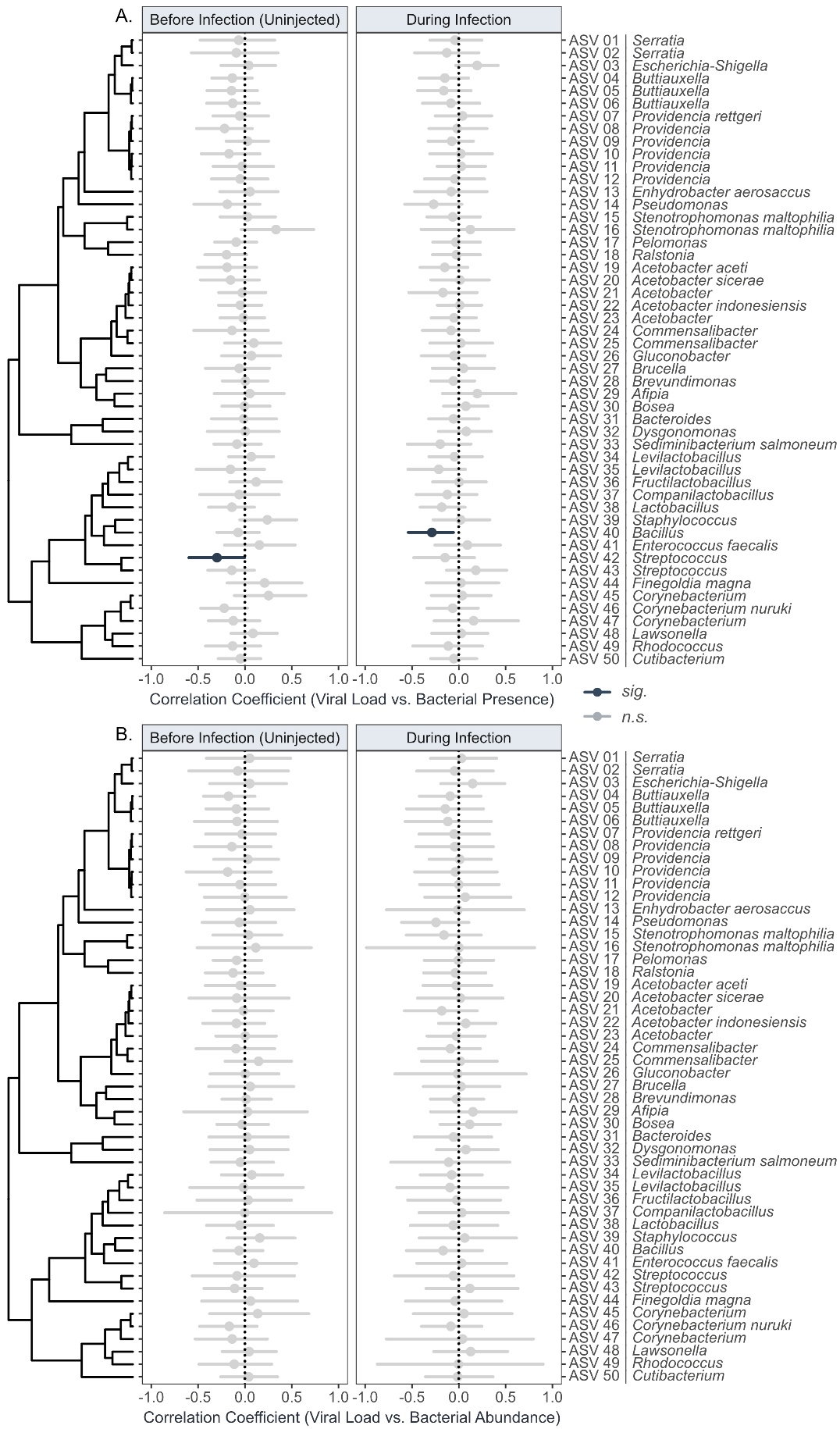

**Supplementary Figure 5: Correlations between viral load and abundance of bacterial ASVs before (Uninjected) and during infection (DCV Infected).** Values for the marginal correlation coefficient (r) between (A) DCV viral load and bacterial presence and (B) Centre Log Ratio (CLR) -transformed abundance using presence and abundances taken from flies before infection (‘uninjected’ condition, left facet) and during infection (right facet). Points represent the mean and bars the 95% HPD intervals of the posterior distribution of each estimate taken from a model of structure (2). Estimates of the correlation coefficient that do not include zero are highlighted in black.
